# An exploration of unusual antimicrobial resistance phenotypes in *Salmonella* Typhi from Blantyre, Malawi reveals the ongoing role of IncHI1 plasmids

**DOI:** 10.1101/2024.08.26.609643

**Authors:** Allan Zuza, Alexander M Wailan, Catherine Anscombe, Nicholas A Feasey, Eva Heinz

## Abstract

Typhoid fever is a significant public health problem endemic in Southeast Asia and Sub-Saharan Africa. Antimicrobial treatment of typhoid is however threatened by the increasing prevalence of antimicrobial resistant (AMR) *S.* Typhi, especially in the globally successful lineage (4.3.1) which has rapidly spread in East and Southern Africa. AMR elements can be found either on plasmids or in one of the three chromosomal integration sites, and there is variability of this across the lineage. Several previous studies with Malawian isolates indicated a clonal, locally spreading lineage with chromosomally integrated resistance genes. In a recent study however we noted three isolates with unusual predicted resistance profiles, and we here present the resolved genomes of these isolates using long- and short-read sequencing. Our work shows that these isolates are potentially imported cases, most closely related to the recently described sub-lineage 4.3.1.EA1, and encode IncHI1 plasmids with reduced resistance profiles compared to the major reference sequence of these plasmids spreading in East Africa. Similar reduced resistance plasmids were reported in a recent large-scale study in five isolates from Tanzania, highlighting the urgency for better coverage of the African continent in genome studies to better understand the dynamics of these potentially co-circulating plasmids.

## INTRODUCTION

*Salmonella enterica* subspecies *enterica* serovar Typhi (*S*. Typhi) is estimated to cause over 11 million typhoid fever cases annually (1). Typhoid caused by *S*. Typhi is endemic in Southeast Asia and Sub-Saharan Africa and a major cause of under-five mortality (1). Treatment of infections by *S*. Typhi relies on antimicrobial treatment; however, the prevalence of *S.* Typhi resistant to first-line antibiotics is rising (2). The spread of antimicrobial resistance (AMR) has been tracked in unprecedented resolution using large-scale whole-genome sequencing surveillance and can clearly be traced to specific lineages that acquire resistance elements and then spread clonally across large areas. Across Africa, there are two main lineages (3); in Eastern Africa the spread of resistant *S.* Typhi is mainly driven by the highly successful lineage 4.3.1 (formerly haplotype H58; (4, 5)) which is often associated with plasmid IncHI1 and which rapidly spread across the African continent from appr. 2005 (6), whilst in Western Africa, the genotypes 2.3.2 and 3.1.1 dominate (3).

The resistance genes commonly found in *S*. Typhi confer resistance to aminopenicillins (*bla*_TEM-1_), chloramphenicol (*catA1*), co-trimoxazole (*dfrA, sul1*/*sul2*), and streptomycin (*strAB*) as well as tetracycline (*tetA*) and can be located either on plasmids or on one of the three chromosomal sites through the integration of a Tn*2670*-like element (3, 6–8), and there is variability of this across the lineage. Previous studies with Malawian isolates have shown the presence of AMR genes in the 4.3.1 haplotype through chromosomal integration (7, 9) and a clonal, locally spreading lineage. The IncHI1 plasmid is known to have substantial variability and is hypothesized to have acquired several resistance gene cassettes through step-wise insertions in an insertion hot-spot (10, 11). Most 4.3.1 isolates in recent large-scale sequencing studies encoding for an IncHI1 plasmid include the core set of resistance genes as described above, which would support the hypothesis that antimicrobial pressure selects for the most resistant lineages which outcompete ones encoding for IncHI1 plasmids encoding for fewer resistances (3, 10).

In a recent study we noted three isolates with unusual resistance profiles; these three were furthermore predicted to carry IncHI1 plasmids which is unusual in the Malawi context (9). We aimed to resolve the genetic structure of these IncHI1 plasmids by performing long-read sequencing and establish whether these were likely imported cases or whether they are part of a lineage spreading in Blantyre carrying the IncHI1 plasmid. To investigate this we selected four representative isolates, including two with unusual resistance profiles for our setting and two as comparisons for putative insertion site changes, and report a detailed description of their genome and plasmid structure, as well as comparisons of the plasmids with reference sequences for the *S*. Typhi IncHI1 plasmid commonly found in 4.3.1.

## METHODS

### Long-fragment DNA extraction

These bacteria were isolated from the blood of febrile patients attending Queen Elizabeth Central Hospital in Blantyre, Malawi, as part of the quality assured routine diagnostic microbiology service supported by the Malawi Liverpool Wellcome Programme as described previously (7). ERS327391 and ERS207185 were previously sequenced by Feasey *et al* and ERS1509723 and ERS1509734 were sequenced by Gauld *et al* (7, 9) using short-read sequencing only. Isolates were recovered from the MLW sample archive and then plated on nutrient agar (Thermo Scientific Oxoid, United Kingdom) and incubated at 37 □ for 18 hours. A single colony was then transferred into 5ml of nutrient broth (Thermo Scientific Oxoid, United Kingdom) and incubated for 18 hours at 37 □. The bacteria cells were then concentrated by centrifugation at 3500rpm for 30 minutes. Total nucleic acids were extracted using the MasterPure complete DNA and RNA Purification Kit (Bioresearch Technologies, United Kingdom) according to the manufacturer’s instructions.

### Sequencing

The double-stranded DNA was quantified using the Qubit 4.0 (ThermoFisher Scientific Inc.) fluorometer and normalized to 400ng in 7.5 ul using UltraPure™ Distilled Water (Invitrogen, Life Technologies Limited). The normalised DNA was used for library preparation using the rapid barcoding kit (SBQ-RBK004, Oxford Nanopore Technologies plc) following the manufacturer’s instructions. The prepared library was sequenced on an Oxford Nanopore r9.4.1 flow cell on the MinION Mk1C sequencer (Oxford Nanopore Technologies plc). Data acquisition was done by the MinKNOW^TM^ software (version 20.10.6, Oxford Nanopore Technologies plc). Base calling and demultiplexing using Guppy Basecalling Software, Oxford Nanopore Technologies plc. (version 6.0.7+c7819bc). We used Guppy with the default settings and the “dna_r9.4.1_450bps_sup.cfg” configuration file for the guppy_barcoder. We used the command line argument “--trim_barcodes” for the debarcoding step using guppy_barcoder. For guppy_barcoder we passed “SQK-RBK004” to the “--barcode_kits” command line argument.

### *De novo* assembly

We retrieved the short reads from ENA and performed trimming and filtering using fastp (version 0.23.1--9f2e255; (12, 13). The reads derived from nanopore sequencing were trimmed and filtered using Filtlong (version 0.2.0--0c4cbe3) with options “--min_length 1000” and “-- keep_percent 90” to discard all reads less than 1kb and remove the worst 10% reads (14). We used FastQC (v0.11.9_cv7) and NanoPlot (version 1.38.1--e303519) with the default settings to check the quality of the filtered short and long reads respectively (15, 16).

We performed long-read-first *de novo* assembly using the Trycycler (v0.5.4) assembly pipeline (17). Briefly, we created 16 read subsamples using Trycycler subsample. We created draft assemblies on each of the simulated reads using either Flye (version 2.9.1-b1780), Minipolish (version 0.1.2) and Raven (version 1.5.0) assemblers (18–20). We used the “--nano-hq” command line argument when generating the Flye assemblies. We run the rest of the assemblers using default settings. We used Any2fasta (version 0.4.2) to convert the gfa formatted output from Minipolish to a fasta file format (21). We used the generated assemblies to create groups of per replicon clusters using trycycler cluster (17). We then performed manual curation steps using trycycler reconcile, trycycler msa, trycycler partition and trycycler consensus following instructions on (https://github.com/rrwick/Trycycler/wiki/How-to-run-Trycycler) (17).

We polished the assemblies using medaka (1.8.1), Polypolish (version 0.5.0) and POLCA (MaSuRCA version 4.1.0) (22–24). The assembly was first polished with the long reads using medaka with “r941_min_sup_g507” passed to the -m command line argument (22). This polished assembly was further polished using short read data using Polypolish and finally using POLCA. Both Polypolish and POLCA were run using default settings (23, 24).

### Genotyping, plasmid calling and AMR gene detection

Bacterial genotyping and plasmid detection was done using the GenoTyphi bacterial typing framework (4, 25). We implemented GenoYyphi using Mykrobe v0.12.1 with the default settings and the Typhi panel (version 20221208; (25, 26)). The json output files were parsed into a csv document using the script “parse_typhi_mykrobe.py” provided at the GenoTyphi repository (26). We then used AMRFinderPlus (Software version 3.11.26 and Database version 2023-11-15.1) to assess the presence of AMR genes and plasmids in the assembled contigs (27). AMRFinder Plus was run by passing the ”–nucleotide –protein –gff –organism –plus” flags. We passed “Salmonella” to the –organism flag. AMRFinderPlus outputs a file with AMR genes and their location on the contigs. The AMR genes are presented in Figure 2. We used ariba (version 2.14.6) using the ARG-ANNOT database to detect AMR genes in the raw reads to annotate the phylogenetic trees (27, 28).

**Figure 1:**
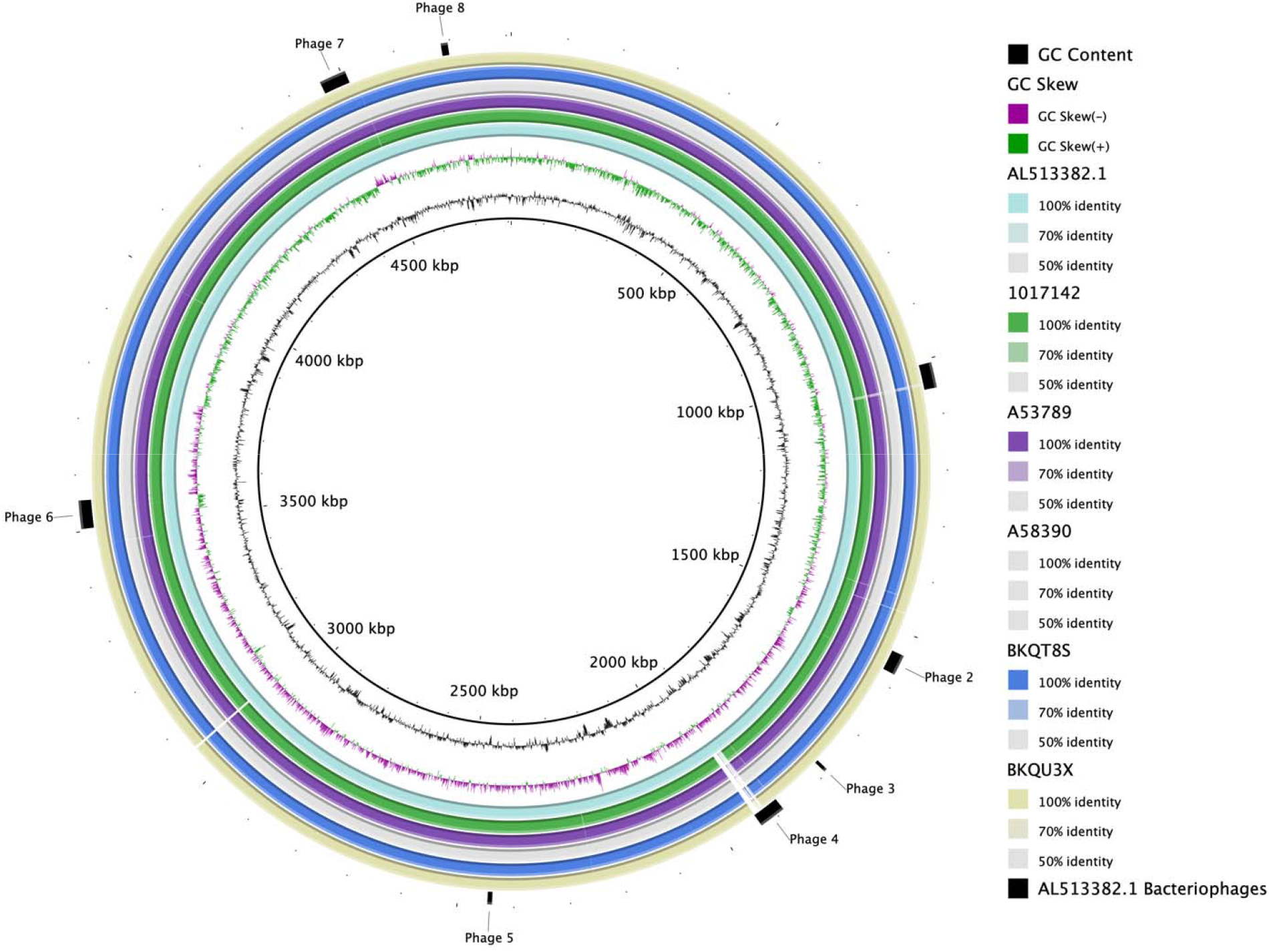
Whole-genome comparison to the CT18 reference. Pairwise BRIG comparison of the hybrid assemblies to the CT18 reference strain as indicated in the legend. The figure also highlights regions containing bacteriophages (outer ring).

**Figure 2:**
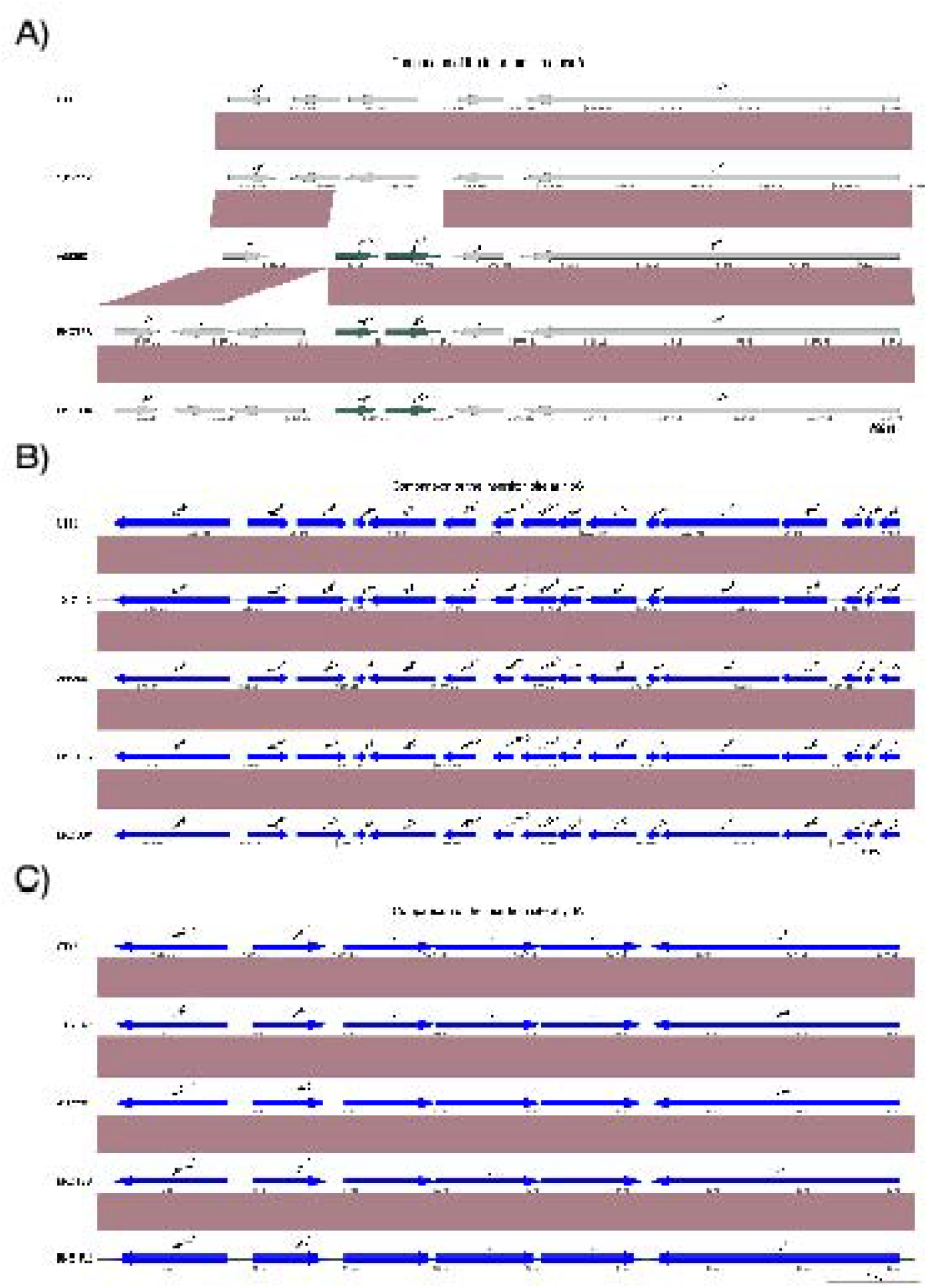
Comparison of the three main described insertion sites for antimicrobial resistance elements. **A)** Comparison of the genomic regions showing the introduction of insertion sequences near the *adenylate cyclase* gene at position 3472059 of the CT18 strain. Conservation of the known insertion sites at positions **B)** 3815480 and **C)** 1690327 of CT18.

### Bacteriophage detection

To detect bacteriophages, we used the PHASTER web tool (https://phaster.ca/; (29, 30)). We loaded the polished assemblies into the web tool and exported the results as a summary.txt file containing the position of phages in the CT18 chromosome, shown in Figure 1 and Figure 2A.

### Annotation

We performed annotation of the hybrid assemblies using Prokka (version 1.14.6) with the default settings and performed a genus-specific annotation with “Salmonella” passed as an argument to the “-–genus” command line option (31).

### Sequence comparison

We used the the Artemis Comparison Tool (ACT release 18.2.0) to perform a comparison of assemblies for the isolates sequenced on the MinION platform (32). We visualised the comparison in the Blast Ring Image Generator (BRIG version 0.95; (33)). Both of these programs used the Basic Local Alignment Tool (BLAST; Nucleotide-Nucleotide BLAST 2.15.0+) to perform nucleotide sequence comparisons (34). For all sequence comparisons, we used the “blastn” option with the default setting for comparing 2 sequences. We used the CT18 reference genome (AL513382.1) as a reference for the BRIG comparison of the chromosome and used the IncHI1 plasmid from the same isolate (AL513383.1) as a reference for the plasmid comparison (35). To assess the presence of AMR genes on the plasmids and parts of the chromosome previously known to contain the insertion of AMR genes, we used the GenoPlotR v 0.8.11 R package which gives a visual comparison of the coding sequences in these regions (34).

### Constructing the phylogenetic tree

We constructed a core snp maximum likelihood phylogenetic tree to put the newly assembled isolates in the context of other isolates from Malawi. The isolates used in the phylogenic analysis are from the paper by Gaud *et al*, where the assembled isolates were first described (7). We used Snippy (version 4.6.0) to perform variant calling and IQ-TREE to construct the phylogenetic tree. We used AL513382.1 as the reference genome for all variant calling steps, snippy-core to create a multiple sequence alignment file with all the variants from the previous step and snp-sites (version 2.5.1) with “-c” argument to extract sites containing exclusively ACGT from the alignment. We also used snp-sites with “-C” to calculate the number of the constant sites which we then used for the “-fconst” argument in IQ-TREE. We performed the IQ-TREE calculation on the alignment with ACGT only generated as described, with the command line parameters “-m GTR+G -bb 1000”.

## RESULTS

### Isolate selection

We performed long-read whole genome sequencing of 4 isolates (ERS327391, ERS207185, ERS1509723, ERS1509734) to better understand unusual antimicrobial resistance patterns in 3 isolates from Malawi as observed previously (9). The years of collection for the isolates were between 2010 and 2016 (9). Isolates ERS1509723 (BKQT8S) and ERS1509734 (BKQU3X) were selected for their AMR pattern, distinct from the rest of the isolates from Malawi from the same period (9). These two isolates are two of only three isolates with IncHI1 plasmids from the same study (9). Isolates ERS327391 (A58390) and ERS207185 (1017142) were selected because they represented isolates at the beginning of the *S*. Typhi outbreak in Blantyre, Malawi (7). Strain A58390 represents one of the first H58 isolates causing illnesses during the *S*. Typhi outbreak. Strain 1017142 represents non-H58 isolates from the same period (7).

### Genome analyses

The nanopore reads for all samples had a mean Phred Quality score of 13.5 with a mean of 29K (21K - 34K) reads per isolate. We generated a mean 275Mb reads (158Mb-338Mb) with a mean N50 of 12K (9k - 13K). We generated hybrid assemblies of the four isolates, accession numbers and their metadata are presented in Table 1. Two isolates (A58390 and 1017142) assembled to 1 contig each, and the other two (BKQT8S and BKQU3X) assembled to 2 contigs each. All isolates had a 4.5Mb contig which represented the chromosome of the isolate. The two isolates with 2 contigs, each had a 185Kb contig that was identified as an IncHI1 plasmid.

**Table 1:**
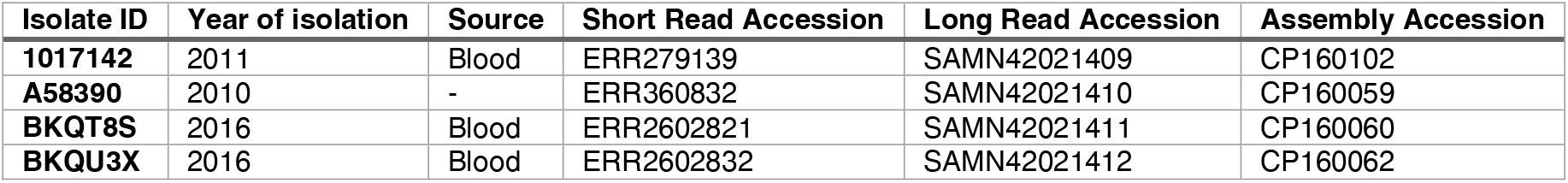
Metadata and accessions for the data presented in this study.

The assembled genomes had a mean GC% of 52.0 (Table 2). Three of the four isolates with long reads belong to the lineage 4.3.1 (H58 haplotype); the two isolates with IncHI1 plasmid were assigned sublineage 4.3.1.1EA1 (36); the remaining isolate belonging to the lineage 4.1.1 (Table 2). A pairwise comparison of the assemblies from this study to the *Salmonella* Typhi reference strain CT18 (AL513382.1) shows a high level of sequence conservation as seen in Figure 1.

**Table 2:** Antimicrobial resistance genes and plasmid replicons.

| Isolate ID | Lineage | Assembly stats |  |  |  | Plasmid replicons |  | AMR determinants |  |  |  |  |  |
| --- | --- | --- | --- | --- | --- | --- | --- | --- | --- | --- | --- | --- | --- |
|  |  | Assembly length | %GC | Contigs | N50 | IncFIAHI1 | IncHI1BR27 | *gyrAS83F* | *bla*<sub>TEM-1D</sub> | *sul2* | *dfrA14* | *tetB* | *aph3*(")-Ib |
| **1017142** | 4.1.1 | 4795235 | 52.05 | 1 | 4795235 | No | No | No | No | No | No | No | No |
| **A58390** | 4.3.1.2 | 4782592 | 52.05 | 1 | 4782592 | No | No | Yes | No | No | No | No | No |
| **BKQT8S** | 4.3.1.1EA1 | 4969415 | 51.82 | 2 | 4783470 | Yes | Yes | No | Yes | Yes | Yes | Yes | Yes |
| **BKQU3X** | 4.3.1.1EA1 | 4969405 | 51.82 | 2 | 4783462 | Yes | Yes | No | Yes | Yes | Yes | Yes | Yes |

We assessed the known chromosomal insertion sites for AMR genes. The insertion site at position 3472059 of the CT18 strain was conserved only in isolate 1017142. The other three isolates had insertion of IS1 insertion sequences (Figure 2A), whilst the insertion sites at 3815480 and 1690327 of the CT18 strain were conserved across the isolates (Figure 2B and C).

### Plasmids, AMR and bacteriophages

The plasmids in the isolates BKQT8S and BKQU3X have an average GC content of 45.94% for each isolate. Both isolates had 3 plasmid replicons of the IncHI1 type (IncFIAHI1, IncHI1A and IncHI1BR27; Table 2) which were all on one plasmid. These multi-replicon plasmids are typed as IncHI1 PST2 using pubMLST (37). A blastn comparison using Megablast shows high levels of similarity between the plasmids (Figure 3A). Comparing the plasmids to the pHCM1 plasmid isolated from the CT18 strain (AL513383.1) indicates a loss of coding sequences in an area with antimicrobial resistance determinants (Figure 3A). The two plasmid sets have a notable difference to pHCM1 in coding sequences in two regions. The first region is located between HCM1.149 and HCM1.175 of the pHCM1 plasmid. This region lost of□17 coding sequences in the plasmids from Malawi. Five of these coding sequences are associated with a mercury resistance operon and one AMR gene (*dhrA14*) has relocated to a different area of the Malawi plasmid. The second region is between HCM1.194 and HCM1.252. This region has been inverted in the Malawian plasmids. The region with the inversion has seen a loss of 13 coding sequences, reducing in size from ∼40kb to ∼20kb. Some of the lost genes include mercury resistance genes and the two AMR genes *catA1* and *aph(6)−Id*. These two AMR genes are responsible for resistance to chloramphenicol and aminoglycosides respectively. Our plasmid maintained aminoglycosides resistance genotype by maintaining a copy of *aph(3’’)−Ib* gene. The plasmid in our collection has managed to maintain its resistance genotype while removing the metal resistance genes which are unnecessary for infection in human hosts. These changes in the structure of the plasmid could indicate adaptation of the plasmid in this host-specific pathogen which could lead to a Typhi population which persistently carries the plasmid without need to relocate the resistance cassette to the chromosome (38). The rearrangements of the AMR genes to a single location allows for centralized control of transcription of these genes, which may lead to better control over the AMR phenotype for isolates carrying this particular plasmid (39). The AMR genes are highlighted in the ring showing the coding sequences of the reference plasmid (Figure 3A). Figure 3B further highlights the region in comparison to the pHCM1 plasmid.

**Figure 3:**
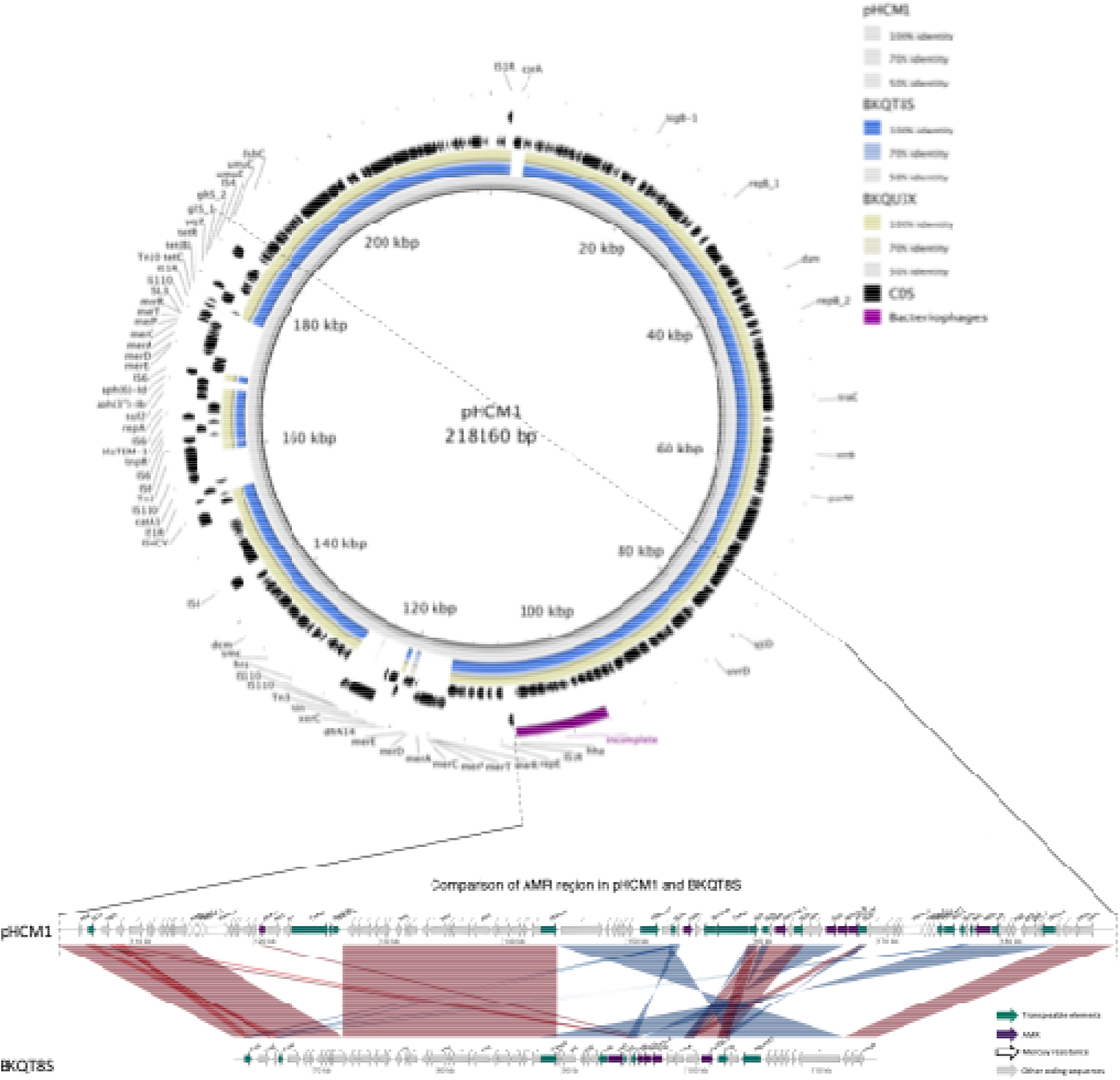
Detailed comparison of the IncHI1 plasmids. **A)** BLAST comparison of the plasmids from this study to the pHCM1 plasmid from the CT18 reference genome of *Salmonella* Typhi. From inside to outside ring: reference sequence pHCM1, plasmid of BKQT8S, plasmid BKQU3X, coding sequences in pHCM1, AMR genes in pHCM1, transposable elements in pHCM1 and bacteriophage region on pHCM1. **B)** BLAST comparison between plasmid of isolate BKQT8S and pHCM1 highlighting the location of IS elements and AMR genes. This region is an expansion of the variable region on Figure 2A. CDS, coding sequences; AMR, antimicrobial resistance.

Neither lineage 4.1.1 isolate, 1017142 nor the 4.3.1 isolate A58390 carried any acquired AMR genes, however there was a *gyrA_S83F* point mutation in A58390 which is associated with reduced susceptibility to fluoroquinolones. This was the only isolate with mutations in the DNA gyrase. The other two isolates each carried acquired resistance genes (*bla*_TEM-1d_, *sul2*, *aph(3”)- lb*, *dfrA14* and *tet(B)*) on the plasmid which are known to cause resistance to beta-lactams, sulfonamides, aminoglycoside, trimethoprim and tetracyclines.

Using PHASTER to define phage elements on the assemblies reveals at least four intact bacteriophages, one putative phage call and several incomplete bacteriophages per isolate Figure 4. Isolates BKQU3X and BKQT8S show similarity in the number of bacteriophage species per region and the completeness of the bacteriophages in these regions (Figure 4). Both isolates contain phages similar in composition and size (Figure 4B).

**Figure 4:**
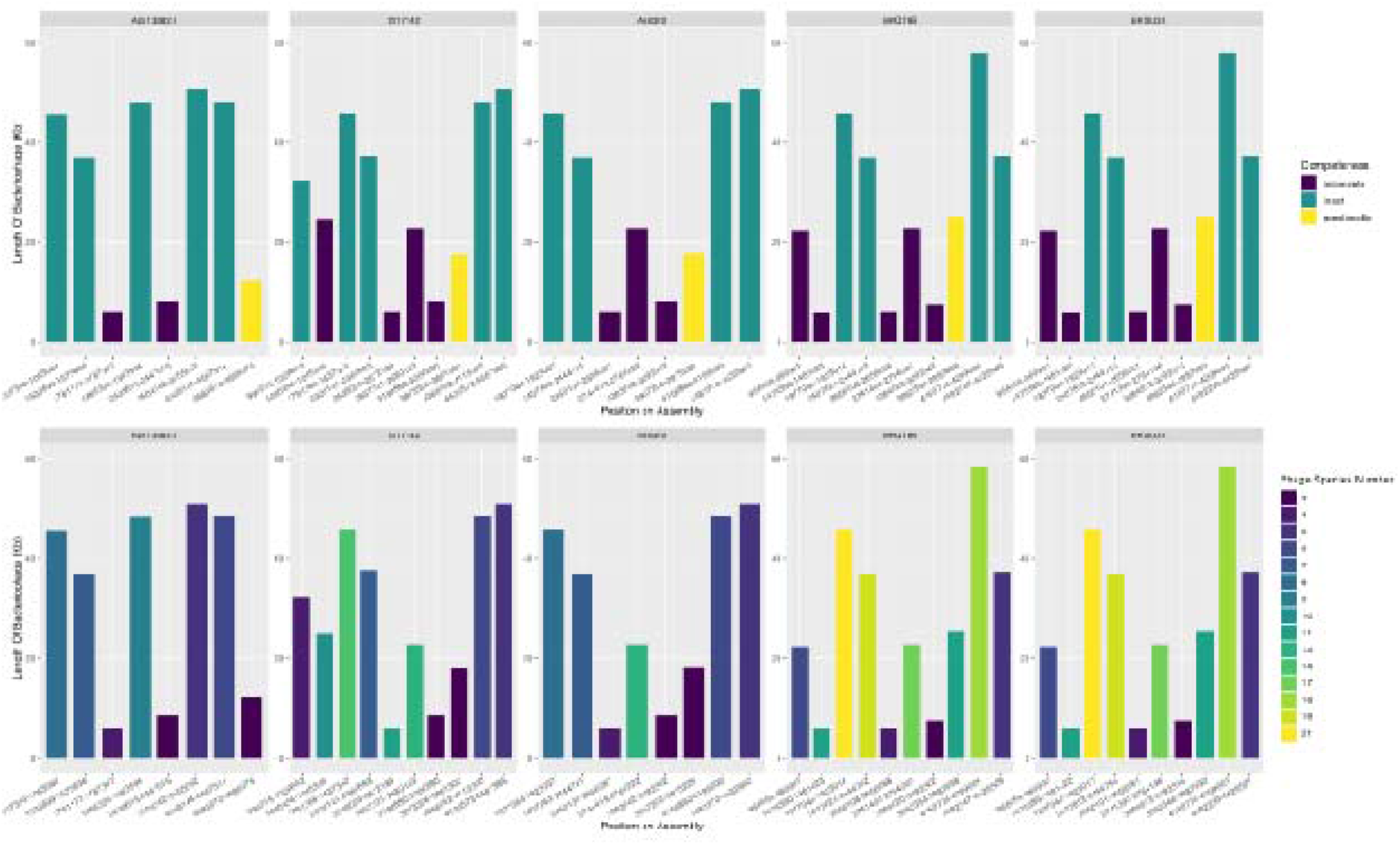
Predicted bacteriophage regions. **A)** Bacteriophage regions on the assemblies. Each column indicates the length of bacteriophage at the position in the CT18 genome colored by the completeness of the bacteriophage at the region**. B)** Each column indicates the length of bacteriophage at the position in the CT18 genome colored by the estimated species numbers (bottom row).

### Phylogenetic analysis

Performing a phylogenetic analysis confirms the 4.3.1.1.EA1 diverse plasmid and AMR profiles (Figure 5). Most of the 4.3.1.1.EA1 isolates do not carry plasmids. Those which carry plasmids seem to be in two plasmid combinations, with isolates from Malawi carrying the plasmid replicon inc_fiahi1 which is missing in the isolates from Kenya, while those from Kenya carry the plasmid replicon inc_hi1_st6 which is not available in the isolates from Malawi. Isolates with plasmids in the Kenyan collection also have additional AMR genes compared to the rest of the 4.3.1.1.EA1 lineage. Isolates which are not 4.3.1.1.EA1 are clustered in distinct clades with 1 clade per lineage. These isolates also have uniform AMR and plasmid profiles. There are distinct differences in acquired AMR genes between isolates of the lineage 4.3.1.1.EA1 and the other H58 lineages. The acquired AMR genes present in the lineage 4.3.1.1.EA1 are absent in the other H58 lineages and vice versa. This might mean that the mobile genetic elements driving resistance for the 4.3.1.1.EA1 are different from those in other H58 lineages.

**Figure 5:**
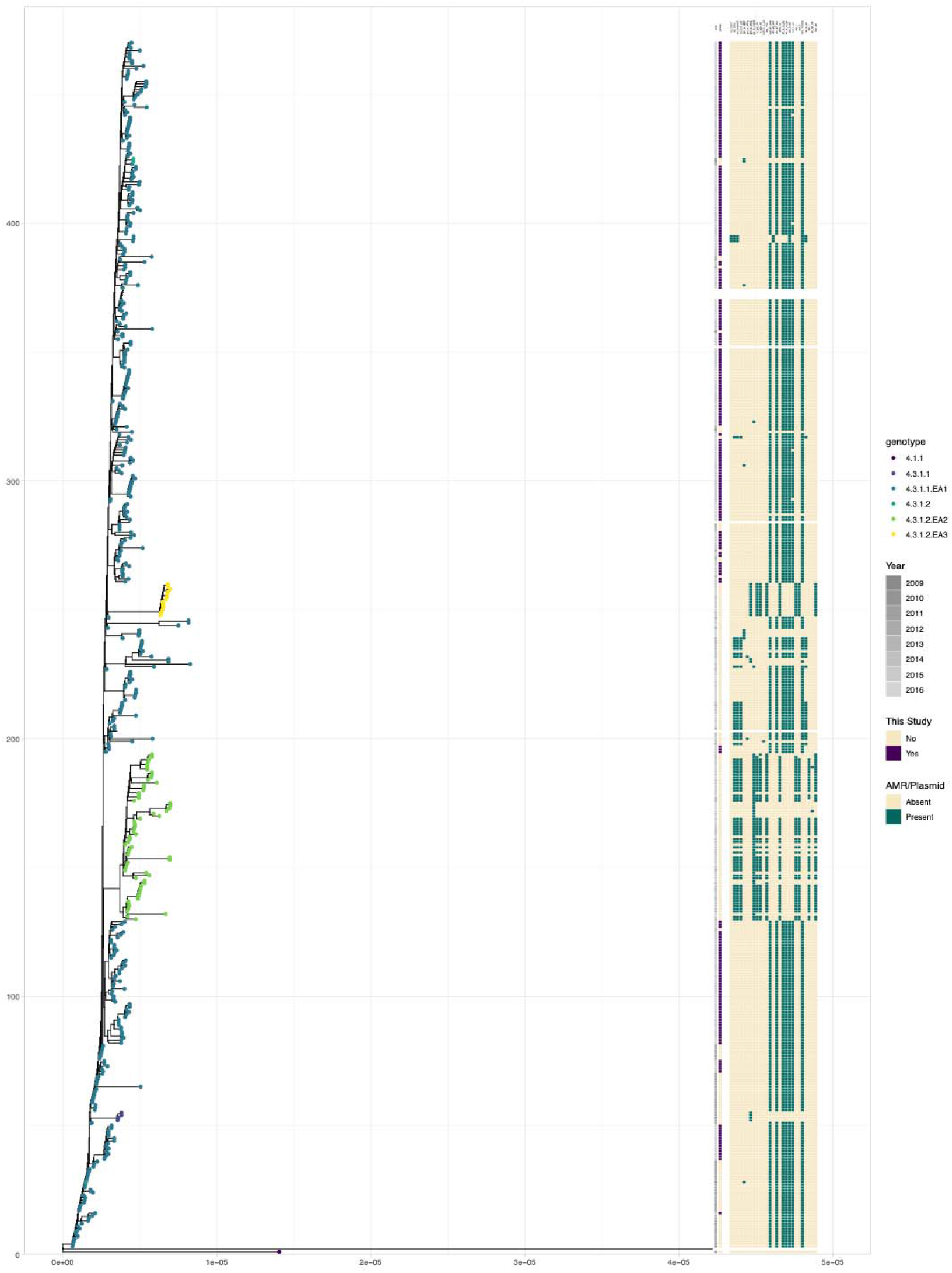
Phylogenetic analysis of the isolates in a wider context. A core-snp maximum likelihood phylogenetic tree of isolates from Malawi into context by including isolates from Kenya from the same period. The tips are coloured by the genotype of the isolate according to the GenoTyphi typing scheme. The bars indicate the year of isolation, and origin of the samples. The remaining bars show the presence of plasmid replicons in the samples, quinolone point mutations and antimicrobial resistance genes in the isolates as identified by ariba, as indicated by the legends.

## DISCUSSION

We noticed three incidences of predicted IncHI1 plasmids present in Malawi *S*. Typhi isolates amongst a set of 335 isolates, with different resistance gene repertoires. These three isolates carry *tetB* and *dfrA14* in addition to *sul2, strA,* and *strB* genes which are found in all other isolates carrying AMR genes, but lack the *cat1*, *sul1* and *tetD* genes usually seen on IncHI plasmids carried by *S*. Typhi. Initial analyses of the assemblies confirmed a single chromosome assembled for the two control isolates, whereas the two isolates with predicted IncHI replicons assembled as two molecules, the chromosome and one plasmid, as expected.

The two unusual isolates were further investigated given their reduced resistance gene repertoire compared to the main IncHI1 plasmid and we identified that these carry the PST2 IncHI1 plasmids, which is not commonly seen in the 4.3.1 haplotype (40). Two studies previously observed PST2 in non-4.3.1 haplotypes among travelers from West Africa to Europe during the same period as our isolates (38, 41). This IncHI variant, also with a reduced set of resistance genes and the plasmid replicons as encoded in our isolates (including a truncated IncFI replicon) were reported initially for the IncHI plasmid R27 in *S*. Typhi (42); an IncFI replicon (although no IncHI replicon) was also noted in several isolates from Tanzania (3).

Initial speculations on *S*. Typhi plasmid diversity considered whether this reduced plasmid might be in the course of being replaced. It is thus interesting to see that, whilst lacking the mercury operons and several resistance determinants, this reduced IncHI plasmid type is extant in parts of south-East Africa and embedded in the highly successful 3.4.1.EA1 lineage. This indicates that selection for the larger resistance plasmid might not be as strong as initially assumed (3) or might even be disadvantageous in specific settings given its larger size.

Our isolates show *S*. Typhi encoding AMR genes either on plasmids or chromosomes isolated from the same region, and we thus investigated the known chromosomal insertion sites in the isolates with IncHI1 plasmids for scars or IS elements (8, 43). We found the insertion of coding sequences and transposases in isolates of the 4.3.1 haplotype. This might suggest the beginning of chromosomal integration of AMR genes previously on the plasmid or represent a scar of lost resistances in this region, although no isolates with acquired AMR genes both on plasmids and the chromosome were identified, which would represent a snapshot of this process in progress. Our data will be a valuable resource for researchers working on comparative genome analyses of *S*. Typhi, in particular in southern Africa, and contribute to improve our understanding of competitive advantages between lineages with different resistance plasmids.

## DATA AVAILABILITY STATEMENT

Raw sequence data is accessible under project ID PRJNA1127853 at SRA and are outlined in Table 1. Assemblies were submitted to GenBank and are accessible under project ID PRJNA1127853, the individual accessions are presented in Table 1.

## AUTHOR CONTRIBUTIONS

Conceptualization: EH, AMW, NAF; Data curation: EH, AZ; Formal analysis: AZ; Funding acquisition: NAF, EH; Investigation: AZ, AMW, EH; Methodology: AZ, EH; Project administration: EH, NAF; Resources: CA, NAF; Software: AZ, EH; Supervision: NAF, EH; Validation: AZ; Visualization: AZ; Writing-original draft: AZ; Writing-review and editing: AZ, NAF, EH. All authors read and approved the final manuscript.

## ACKNOWLEDGMENTS

We want to thank the MLW core informatics support team and Philip M Ashton for expert technical support.

